# Chromatin modifiers and recombination factors promote a telomere fold-back structure, that is lost during replicative senescence

**DOI:** 10.1101/2020.01.13.904086

**Authors:** Tina Wagner, Lara Perez-Martinez, René Schellhaas, Marta Barrientos-Moreno, Merve Öztürk, Félix Prado, Falk Butter, Brian Luke

## Abstract

Telomeres adopt a lariat conformation and hence, engage in long and short distance intra-chromosome interactions. Budding yeast telomeres were proposed to fold back into subtelomeric regions, but a robust assay to quantitatively characterize this structure has been lacking. Therefore, it is not well understood how the interactions between telomeres and non-telomeric regions are established and regulated. We employ a telomeric chromosome conformation capture (Telo-3C) approach to directly analyze telomere folding and its maintenance in *S. cerevisiae*. We identify the histone modifiers Sir2, Sin3 and Set2 as critical regulators for telomere folding, which suggests that a distinct telomeric chromatin environment is a major requirement for the folding of yeast telomeres. We demonstrate that telomeres are not folded when cells enter replicative senescence, which occurs independently of short telomere length. Indeed, Sir2, Sin3 and Set2 protein levels are decreased during senescence and their absence may thereby prevent telomere folding. Additionally, we show that the homologous recombination machinery, including the Rad51 and Rad52 proteins, as well as the checkpoint component Rad53 are essential for establishing the telomere fold-back structure. This study outlines a method to interrogate telomere-subtelomere interactions at a single unmodified yeast telomere. Using this method, we provide insights into how the spatial arrangement of the chromosome end structure is established and demonstrate that telomere folding is compromised throughout replicative senescence.

**Author summary:** Telomeres are the protective caps of chromosome ends and prevent the activation of a local DNA damage response. In many organisms, telomeres engage in a loop-like structure which may provide an additional layer of end protection. As we still lack insight into the regulation of the folded telomere structure, we used budding yeast to establish a method to measure telomere folding and then study the genetic requirements for its establishment. We found that cells require the homologous recombination machinery as well as components of the DNA damage checkpoint to successfully establish a folded telomere. Through the deletion of telomerase in budding yeast, we investigated how telomere folding was regulated during replicative senescence, a process that occurs in the majority of telomerase negative human cells. During senescence, telomeres gradually shorten and erode until cells stop dividing which is a potent tumor suppressor and prevents unscheduled growth of potential cancer cells. We found, that the folded telomere structure is compromised as part of the cellular senescence response, but not due to telomere shortening *per se.* We think, that an altered telomeric chromatin environment during senescence is important to maintain an open state – which may be important for signaling or for repair.

## Introduction

Telomeres are essential nucleoprotein structures at the physical ends of eukaryotic chromosomes consisting of non-protein coding DNA repeats and telomere bound protein complexes. In human cells, telomere structure and function are controlled by a six-protein complex called Shelterin, that binds to telomeres in a sequence specific manner (1). In yeast, these functions are executed by the CST (Cdc13-Stn1-Ten1) complex (2) together with Rap1, Rif1 and Rif2 (3–5). Although telomeres resemble a one-ended DNA double strand break (DSB) they are refractory to being acted upon by the DNA damage response (DDR) (6). Hence, telomeres prevent illegitimate repair events that would cause chromosome end-to-end fusions and lead to genome instability (6). This “end protection” property of telomeres has largely been attributed to the associated protein binding complexes, but a telomeric loop structure (t-loop) also appears to be critical. Telomeric lariat structures have been demonstrated in species ranging from yeast to human (7–13), however, their properties may vary considerably between organisms. In mammalian cells, the t-loop forms within the telomeric repeat tract, via strand-invasion of the 3’ overhang into the double stranded region of the telomere (13, 14). In contrast, telomeres in *S. cerevisiae* are proposed to fold back into the subtelomeric region, where they are maintained by an, as of yet, unknown mechanism (12,15,16).

Little information exists about the regulation of telomere fold-back structures. TRF2, a component of the Shelterin complex, is required for t-loop formation in human cells (13, 14). In addition to promoting t-loops, TRF2 also regulates their dissolution in S-phase via recruitment of the helicase RTEL1 (17, 18). In *S. pombe* the telomeric protein Taz1 has been demonstrated to remodel telomeric DNA into loops (8) and in *S. cerevisiae* histone deacetylases have been shown to be required for the fold-back into the subtelomere (15, 19). In addition, a genome-wide screen using a fold-back reporter gene on a modified telomere in mutants of the non-essential yeast deletion collection, indicates that other protein families, including chromatin remodelers and transcription regulators, are also involved in the maintenance of telomere structure (15).

Human and mouse t-loops can be visualized via electron- and super resolution-microscopy (13,14,20). The existence of looped structures could also be shown microscopically in *K. lactis* harboring overelongated telomeres (11). The short length, and base composition, of *S. cerevisiae* telomeres prevent such microscopy-based approaches, hence only genetic- and chromatin immunoprecipitation (ChIP) - based experiments have been employed to study telomere folding in yeast (12,15,16,19). A major concern of the genetic approach is that the subtelomere is deleted and modified with a reporter (12,15,19), which may strongly alter protein composition and chromatin status in the vicinity of the telomere. On the other hand, ChIP of telomeric proteins to the subtelomere may indeed represent telomere folding, however the spreading of telomere binding proteins into subtelomeric regions cannot be completely ruled out (15, 16).

Telomeres become shortened with each cell division due to the end replication problem (21–23). The loss of telomeric repeats leads to the progressive loss of chromosome end protection and replicative senescence, a permanent cell cycle arrest. To counteract telomere shortening, certain mammalian cell types, yeast cells and >85 % of cancer cells express the reverse transcriptase telomerase, which adds telomeric sequences to the 3’ end of the shortest telomeres (24–26). Yeast constitutively express telomerase but deletion of the catalytic subunit (*EST2*) or the RNA component (*TLC1*) of telomerase renders them susceptible to the end replication problem and eventually replicative senescence (26). As a rare event, some cells (survivors) can overcome senescence and crisis by elongating their telomeres via homologous recombination-based mechanisms (27).

Critically short telomeres arising during replicative senescence activate the DNA damage checkpoint and are recognized by DNA repair factors, predominantly by the protein kinase Mec1 (ATR) but also by Tel1 (ATM), both master regulators of the DSB response (28–31). The activation of either kinase triggers the phosphorylation of the effector kinases Rad53 and Chk1, which leads to a cell cycle arrest. Additionally, the homology directed repair (HRD) proteins Rad51 and Rad52 co-localizes with critically short telomere in the absence of telomerase to promote their extension (repair) (32, 33). Surprisingly, it was shown that proteins involved in DDR, such as the MRX (Mre11, Rad50, Xrs2) complex and Tel1 also localize to telomeres in telomerase positive cells and play a general role for telomere stability and telomere length regulation (34). Cells that lack both Mec1 and Tel1 show a progressive telomere shortening and undergo replicative senescence (35, 36) in addition to harboring increased telomere fusions (37–40). At mammalian telomeres, the DNA damage response and the homologous recombination machinery also play a role at functional telomeres. It was reported for human cells, that ATM and ATR are transiently recruited to telomeres during telomere replication where they activate a local DDR. (41, 42). After telomere processing is complete, Rad51 and Rad52 are recruited and likely promote strand invasion of the 3’ overhang into the double-stranded telomeric repeats to generate a t-loop. This transient localization does not result in unscheduled recombination and sister chromatid exchange between telomeres. Recent studies have indicated that the t-loop structure plays an important role in the regulation of the DNA damage response at telomeres (20, 43). When the t-loop is not formed, telomeres aberrantly activate an ATM response, however they remain resistant to telomere fusions.

Here, we developed telomeric chromosome conformation capture (Telo-3C) at a natural yeast telomere, which provides a direct readout for the telomere folding structure into the subtelomere. Wild type length telomeres are able to fold-back (and form a “closed state”). However, when the HDR proteins Rad51 or Rad52 are not present, telomere folding occurs less frequently, suggesting a strand invasion mechanism similar to what has been described for the t-loop in human cells. Consistent with the DDR being involved in fold-back establishment, telomere folding is also defective when the checkpoint kinase, Rad53, is depleted. As cells enter replicative senescence, we show that telomere folding is not maintained. We conclude that this “open state” is not a result of telomere shortening *per se*, but seems to be linked to the senescence program. We identified the histone modifiers Sir2, Sin3 and Set2 as potential regulators of telomere folding during replicative senescence. Indeed, cells have a decreased capacity to establish telomere folding when lacking any of these factors. We hypothesize that the reduced expression of these factors likely contributes to ensuring an open telomere state during senescence. Taken together, out data offer new insights into understanding how telomeres physically interact with other regions of the genome.

## Results

### Telo-3C detects telomere folding in *S. cerevisiae*

To gain insights into the dynamic regulation of telomere folding in *S. cerevisiae*, we developed a telomere chromosome conformation capture (3C) approach (44) (Telo-3C). This provides a direct readout to quantitatively assess the interaction of one telomere (left telomere of chromosome 1) with its adjacent subtelomeric region. As illustrated in Fig 1A, chromosome contacts were crosslinked *in vivo* using formaldehyde and the extracted chromatin was digested with a suitable restriction enzyme (NcoI for telomere 1L) (see Fig 1B for precise restriction sites). The compatible DNA ends were ligated under very dilute conditions, resulting in the ligation of only those loci that were in close proximity at the time of the crosslink. After reversing the crosslink, a specific 3C ligation product can be amplified from primers that were initially co-directionally oriented on genomic DNA (no PCR product) but get converted through the 3C reaction to being convergent (giving a PCR product, see blue arrows Fig 1A).

**Figure 1:**
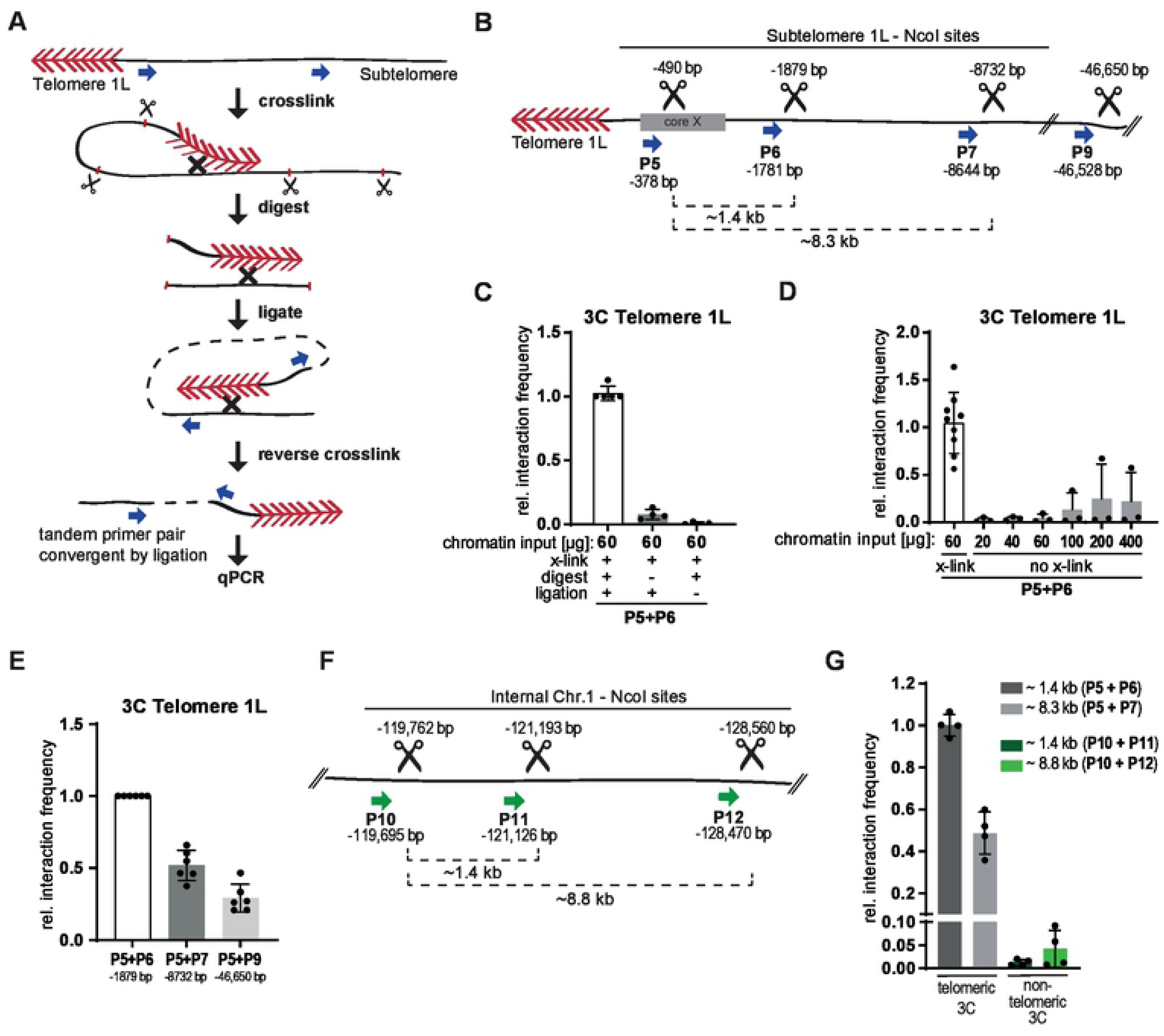
Telo-3C detects telomere folding in *S. cerevisiae*. **A.** Schematic representation of the Telo-3C method. Formaldehyde was used to crosslink chromosome interactions such as telomere folding. DNA was digested with the restriction enzyme NcoI and DNA ends were ligated under diluted DNA conditions. Previously tandem primer pairs (blue arrows) got converted to convergent primer pairs by ligation, and telomere folding was detected by qPCR. Red arrowheads = telomere repeats; Scissors = NcoI restriction sites **B.** Illustration of NcoI cut sites along the subtelomere 1L and the primers (blue arrows) used to detect interaction frequencies between the respective sites. P5 was used as anchor primer in close proximity to the telomeric repeats. Genomic location of primers and cut sites (scissors) were calculated from the end of telomere 1L (depicted in bp) as annotated on SGD (*Saccharomyces* genome database). **C.** The Telo-3C protocol was performed without digesting or ligating the chromatin as indicated. The interaction frequency was normalized to an intergenic PCR product. The rel. interaction frequency of the sample which was performed with the complete Telo-3C protocol was set to 1. Technical replicates were performed for this sample. Mean +/- SD of 4 independent experiments. **D.** The Telo-3C protocol was performed on increasing amounts of chromatin which was not crosslinked (x-link) with formaldehyde. The interaction frequency was normalized to an intergenic PCR product. The rel. interaction frequency of the sample which was performed with the complete Telo-3C protocol was set to 1. Mean +/- SD of 3 independent experiments. Technical replicates were performed for the non-x-linked samples. **E.** Relative interaction frequencies of telomere 1L with loci of increasing distance from the telomeric repeats (primers P6, P7 and P9). The interaction frequency was normalized to an intergenic PCR product. The rel. interaction frequency of P5 and P6 was set to 1. Mean +/- SD of 6 independent experiments. **F.** Illustration of NcoI cut sites along chromosome 1 (non-telomeric) and the primers (green arrows) used to detect interaction frequencies between the respective sites. Genomic location of primers and cut sites were calculated from the end of telomere 1L as annotated on SGD. **G.** Telo-3C results comparing the relative interaction frequencies of telomeric with non-telomeric loci on chromosome 1 using the indicated primer pairs. The interaction frequency was normalized to an intergenic PCR product. The rel. interaction frequency of P5 and P6 was set to 1. Mean +/- SD of 4 independent experiments.

To ensure the specificity of the Telo-3C procedure we performed the protocol without ligating or digesting the chromatin. In these conditions, we did not detect a 3C product of the same intensity as compared to the complete Telo-3C protocol (Fig 1C). Similarly, when the chromatin was not crosslinked with formaldehyde, a 3C PCR product was not detectable even with increasing amounts of chromatin input (Fig 1D). This suggests that a specific 3C PCR product could only be amplified when the loci of interest were located in close proximity. We were interested in measuring the interaction of the telomeric repeats with several loci along the subtelomere and designed primers according to the NcoI restriction sites that were available (Fig 1B). Primer P5, directly juxtaposed to the telomere repeat sequences, was used as an anchor primer in combination with P6, P7 and P9 thereby comparing the interaction of the telomeric repeats to a subtelomeric region with a distance of ∼2 kb (P6) and ∼9 kb (P7). With P9 we chose a primer outside the region which we would define as subtelomere with a distance of ∼47 kb from the telomeric repeats. The interaction frequency was reduced to half when moving from a ∼2 kb (P6) to ∼ 9 kb (P7) distance from the telomeric repeats (Fig 1E). We also detected a 3C product between primers P5 and P9 (∼47 kb) which suggests that the telomere may also interact with loci of greater distance to the telomeric repeats, although with a reduced frequency (Fig 1E). The sequence of all depicted 3C PCR products was confirmed by Sanger sequencing. In order to understand if telomere folding occurs more frequently than a random interaction of two loci on the same chromosome, we analyzed chromosome interactions at internal NcoI restriction sites on chromosome 1 (Fig 1F). The respective 3C primers (P10 (anchor), P11 and P12) have similar distances as the ones used on subtelomere 1L. Although we could detect a weak interaction between loci using the internal primer sets, the interactions between telomere 1L and its subtelomere were more robust. These results demonstrate that the interaction at the end of the chromosome occurs more frequently than a random encounter of two loci on the same chromosome (Fig 1G).

In summary, we established and validated a 3C method that detects the direct interaction of telomeric and subtelomeric regions on a single chromosome, which provides further evidence for the presence of a folded structure at chromosome ends in budding yeast. It appears, that telomere folding occurs more frequently, or more stably, than internal interactions on the same chromosome. Moreover, the telomeric repeats might touch down at multiple regions, and the interaction frequency diminishes with increasing distance from the telomeric tract.

### Telomere 1L opens during replicative senescence

To understand how telomere structure may be regulated throughout the process of replicative senescence, we passaged budding yeast lacking the RNA component of telomerase (*TLC1*) over multiple generations to induce gradual telomere shortening (45). As expected, the *tlc1* cells progressively experienced decreased growth potential (Fig 2A). The loss of viability was caused by telomere shortening due to the lack telomerase activity, as shown in a telomere restriction fragment analysis (Fig 2B, representative example of one clone). At crisis, the point of lowest viability, some cells can overcome the permanent cells cycle arrest and have acquired the capacity to elongate their telomeres based on a recombination-mediated pathway (27). These survivors showed the amplification of TG repeats by the appearance of multiple additional bands on the Southern blot (46) (Fig 2B). We employed Telo-3C during this time course and measured the interaction of the telomere 1L with its subtelomere at the indicated time points (+ in Fig 2A). Telo-3C analysis at PD 9 revealed that telomere folding of *tlc1* mutants was comparable to wt telomeres, despite their telomeres being much shorter that wt telomeres (Fig 2B, C). However, when *tlc1* cells had critically short telomeres and approached the point of telomeric crisis, telomere folding decreased significantly (PD 47, 56 and 64 in Fig 2C). In survivors, however (PD 101 in Fig. 2C), telomeres re-acquired the ability to establish a fold-back structure. Using a probe specific for telomere 1L, we confirmed that the length of telomere 1L approximately followed the same shortening and recombination pattern as bulk telomeres (S1A Fig). Flow cytometry analysis for DNA content confirmed that the analyzed cell populations did not demonstrate drastic changes in terms of cell cycle distribution (S1B Fig).

**Figure 2:**
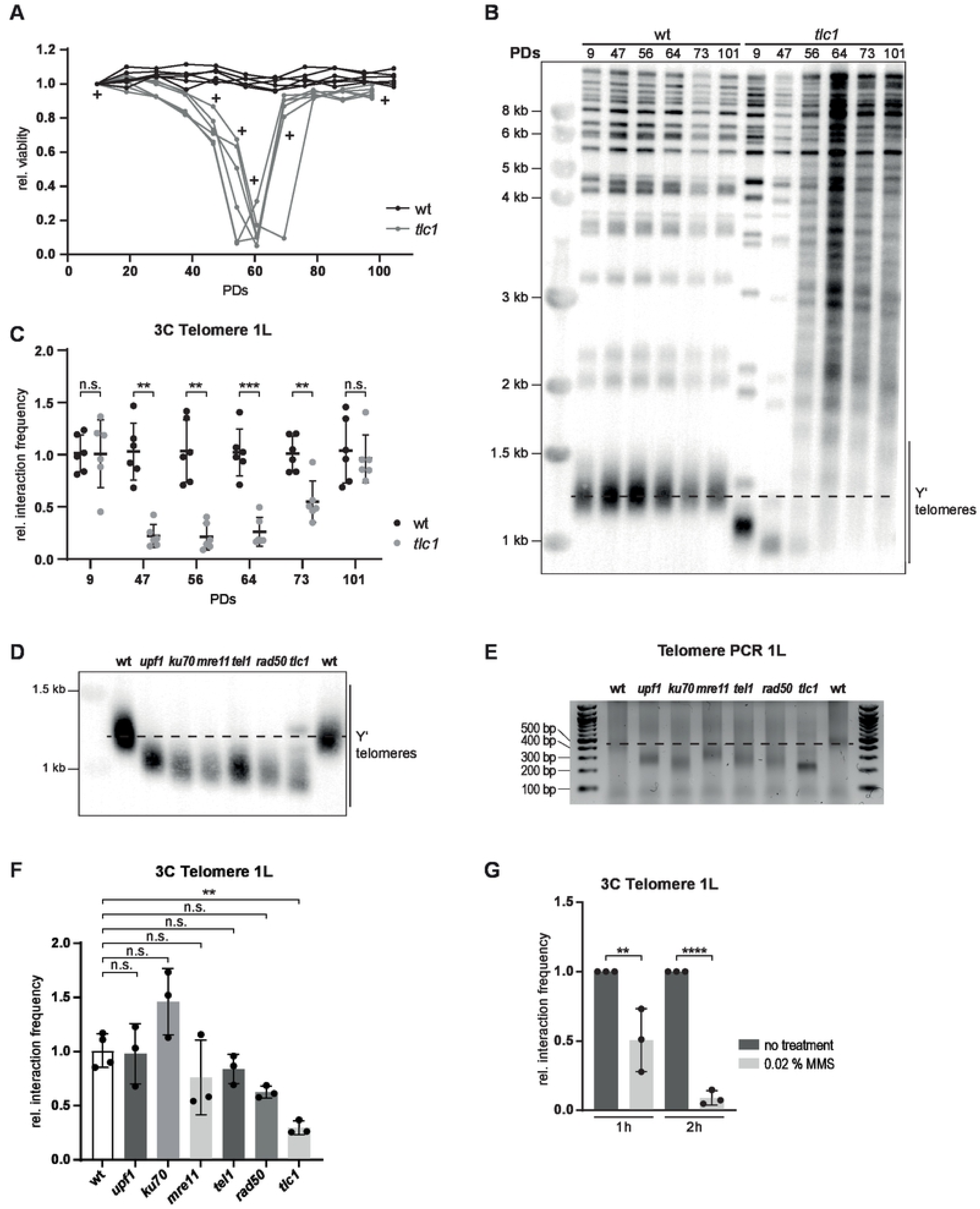
Telomere 1L opens during replicative senescence. **A.** The relative viability of 6 independent wt and *tlc1* cultures was followed over 100 PDs. Samples for Telo-3C were collected at the indicated time points (+). We define PD0 as the time the senescence curve was started in liquid media from the germinated spore. **B.** Southern blot for all telomeres on XhoI digested genomic DNA using a radio-labeled (TG)n-repeat containing vector fragment. One representative sample for each genotype is shown. The dotted line indicates wt telomere length. **C.** Telo-3C analysis during the time course shown in (A). The relative interaction between primers P5 and P6 (Fig 1B) is shown. The interaction frequency was normalized to an intergenic PCR product. The rel. interaction frequencies were normalized to the wt of the respective day. Mean +/- SD of 3 independent experiments, which 2 technical replicates each. Adjusted p-values were obtained from two-way ANOVA with Sidak’s multiple comparison test. **D.** Southern blot for all telomeres on XhoI digested genomic DNA from the indicated deletion mutants using a radio-labeled (TG)n-repeat containing vector fragment. The *tlc1* mutant was analyzed at PD40 as calculated in Fig 2A. A representative sample for each genotype is shown. The dotted line indicates wt telomere length. **E.** Telomere PCR for the length of telomere 1L in the depicted mutants from the genomic DNA used in Fig 2D. Dashed red line indicates the wt telomere length. **F.** Telo-3C analysis of the displayed mutants. The interaction between primers P5 and P6 is shown. The interaction frequency was normalized to an intergenic PCR product. The rel. interaction frequency of wt was set to 1. The *tlc1* mutant was analyzed at PD40 as calculated in 2A. Mean +/- SD of 3 independent experiments. Technical replicates were performed for the wt. Adjusted p-values were obtained from one-way ANOVA with Dunnett’s multiple comparison test. **G.** Telo-3C analysis of wt cells after 1 h and 2 h of MMS treatment (0.02 %). The interaction between primers P5 and P6 is shown. The interaction frequency was normalized to an intergenic PCR product. The rel. interaction frequency of the untreated sample was set to 1. Mean +/- SD of 3 independent experiments. Adjusted p-values were obtained from two-way ANOVA with Sidak’s multiple comparison test. PDs = population doublings. (*p ≤ 0.05, **p ≤ 0.01, ***p ≤ 0.001, n.s., not significant)

The fact that telomeres remain folded at PD 9 (Fig 2C) suggests that telomere shortening alone is not responsible for loss of the folded structure. To address this further, we employed Telo-3C in several yeast mutants with very short, albeit stable, telomeres. All of the mutants analyzed harbored telomeres (both bulk telomeres and 1L specifically) that were comparable in length to telomerase negative *tlc1* cells but showed neither progressive shortening nor a senescence phenotype (Fig 2D, E). We found that the telomere 1L of these mutants established a fold-back structure that varied in strength between mutants, but was consistently more frequent than in *tlc1* mutants (Fig 2F). This suggests that short telomeres *per se* are still able to fold back onto the subtelomere.

As replicative senescence is induced by a cell cycle arrest due to a DNA damage response at telomeres, we investigated whether other forms of DNA damage would also result in an open telomere state. Upon treatment of wild type cells with the DNA damaging agent methyl methanesulfonate (MMS), we observed decreased telomere folding, that became more pronounced over time (Fig 2G).

In conclusion, we demonstrate that critically short telomeres open their folded structure during replicative senescence and prolonged exposure to MMS. The opening of the fold-back structure is not solely due to the short nature of these telomeres during senescence. Instead, the decreased telomere folding ability during crisis might rather be due to other features of the “senescence program” of these cells.

### Rad51, Rad52 and Rad53 are required for telomere folding

In order to identify regulators of telomere folding, we first employed a candidate approach of factors that could potentially influence telomere fold-back formation and maintenance. First, we chose to investigate the impact of the structural maintenance of chromosomes (SMC) complexes on telomere folding. Cohesin and condensin have been demonstrated to play important roles in terms of chromosomal architecture, including chromatin loop formation and chromatin condensation, respectively (47). We also wanted to test if the SMC5/6 complex might be involved in telomere folding due to its described roles at telomeres (48–50). We tagged one subunit of each SMC complex with an Auxin-inducible degron (AID*) (Morawska and Ulrich 2013) (Fig 3A). As all of these complexes are essential in yeast, this system allowed us to study their impact on telomere folding by degrading the tagged protein through the addition of the plant hormone auxin (indole-3 acetic acid = IAA) to the culture media. We treated the culture with auxin for 1 h, which resulted in a nearly complete depletion of the respective proteins (Fig 3A), while not impacting the viability of the cells (S2A Fig). Telo-3C results showed, that telomere folding was not affected by the lack of any of the SMC complexes under these conditions (Fig 3B). Therefore, although the SMC complexes are implicated in chromosome compaction (51–53) and the establishment of chromosome domain boundaries and chromatin loops (54, 55), they are not a requirement for telomere folding.

**Figure 3:**
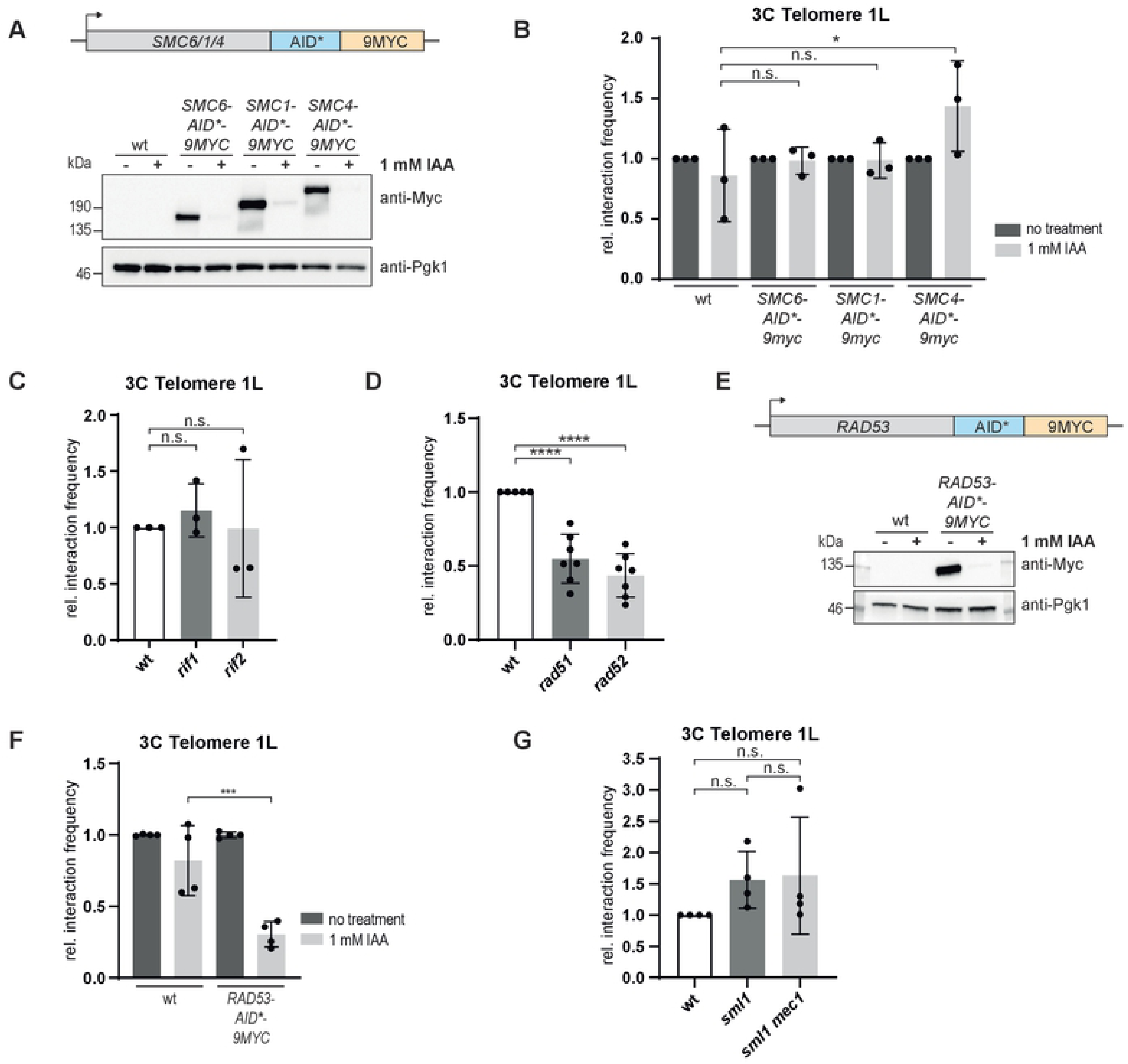
Rad51, Rad52 and Rad53 are required for telomere folding. **A.** *SMC6*, *SMC*1 and *SMC4* were tagged with an auxin-inducible degron (AID*) and a 9xMYC-tag. Western blots were performed to detect protein levels after treatment with 1 mM IAA for 1 h. **B.** Telo-3C analysis after depletion of SMC complex members for 1 h with 1 mM IAA. Mean +/- SD of 3 independent experiments. Adjusted p-values were obtained from two-way ANOVA with Tukey’s multiple comparison test. **C.** Telo-3C analysis of *rif1* and *rif2* mutants. Mean +/- SD of 3 independent experiments. Adjusted p-values were obtained from one-way ANOVA with Dunnett’s multiple comparison test. **D.** Telo-3C analysis of *rad51* and *rad52* mutants. Mean +/- SD of 5 independent experiments. Technical replicates were performed for the mutants. Adjusted p-values were obtained from one-way ANOVA with Dunnett’s multiple comparison test. **E.** *RAD53* was tagged with an auxin-inducible degron (AID*) and a 9xMYC-tag. Western blot to detect protein levels after treatment with 1 mM IAA for 2 h. **F.** Telo-3C analysis after depletion of Rad53 for 2 h with 1 mM IAA. Mean +/- SD of 3 independent experiments, including technical replicates. Adjusted p-values were obtained from two-way ANOVA with Tukey’s multiple comparison test. **G.** Telo-3C analysis of *sml1* and *sml1 mec1* mutants. Mean +/- SD of 4 independent experiments. Adjusted p-values were obtained from one-way ANOVA with Tukey’s multiple comparison test. For all Telo-3C results, the interaction between primers P5 and P6 is shown. The interaction frequency was normalized to an intergenic PCR product. The rel. interaction frequency of the untreated sample from each genotype or the respective wt sample was set to 1. IAA = Indole-3 acetic acid. (*p ≤ 0.05, **p ≤ 0.01, ***p ≤ 0.001, n.s., not significant)

We tested the telomeric proteins Rif1 and Rif2 for their contribution to establishing telomere folding. Both Rif1 and Rif2 were previously reported to regulate telomere structure as measured by the fold-back reporter gene (15). However, by Telo-3C we did not observe a telomere folding defect in either *rif1* or *rif2* deletion mutants arguing that these two telomeric proteins might not contribute to the establishment of telomere folding (Fig 3C). This discrepancy between the two methods might arise from the two different readouts employed, or due to the introduction of an artificial subtelomere with the reporter gene. Additionally, the chromatin structure at telomeres of *rif1* and *rif2* mutants is altered which affect natural and modified subtelomeres in different ways (4, 5).

In human cells the t-loop consists of a D-loop recombination intermediate, which is suggestive of a strand invasion event. To test whether recombination may also be involved in yeast telomere folding, we performed Telo-3C experiments in cells lacking the central HDR factors *RAD51* and *RAD52*. We found that telomere folding was decreased in both *rad51* and *rad52* mutants (Fig 3D). This strongly suggests that in yeast, like in human cells, a strand invasion contributes to the establishment of a closed telomere state.

As Rad51 and Rad52 promote repair in response to a DDR, we tested whether the most upstream factors of the DDR, the checkpoint kinases Mec1 (ATR) and Rad53 (CHK2), may also contribute to telomere folding. We depleted cells of Rad53 for 2 h using an auxin-inducible degron (Fig 3E), a treatment which did not result in decreased cell viability (S2B Fig). Under these conditions, we observed a significant reduction of telomere folding (Fig 3F) suggesting that an impaired DNA damage checkpoint does affect the structure of the telomere. Surprisingly, the deletion of either *MEC1* (Fig 3G) or *TEL1* (Fig 2F) alone did not affect the folding behavior of telomere 1L. We propose, that Mec1 and Tel1 might have overlapping functions in terms of telomere folding which could potentially compensate for each other. Testing the *tel1 mec1* double mutant is complicated by the fact that it induces replicative senescence (35) and a has high incidence of telomere fusions (37, 39). Alternatively, Rad53 may promote telomere folding in a Mec1 and Tel1 independent manner.

### Loss of Sir2, Sin3 and Set2 contribute to telomere opening in senescence

In order to understand the underlying mechanisms behind the structural changes that occur at telomeres throughout senescence, we analyzed the whole proteome of *tlc1* cells as they approached telomeric crisis and compared it to wt cells. Label-free quantitative proteomics analysis revealed 751 significantly downregulated and 678 upregulated proteins during senescence (Fig 4A, see S3 Fig for a comprehensive list and S4A, B Fig for gene ontology analysis). To identify candidates that potentially regulate telomere folding during senescence, we compared results from a previous screen for regulators of telomere folding in *S. cerevisiae* (15) with downregulated proteins from our proteomics data set. This genetic screen by Poschke et al., identified 113 genes and we manually added *sir2* to this list. Sir2 has been demonstrated to promote telomere folding however its deletion cannot be detected in the screening procedure due to technical reasons (12,19,56). We found 21 overlapping candidates between the fold-back defective mutants and proteins that become downregulated during senescence (Fig 4B). Downregulation of these candidate proteins during senescence may therefore contribute to the loss of a closed telomere state. Compared to the complete set of downregulated proteins during senescence, the 21 candidates (Fig 4C) were significantly enriched for the gene ontology terms of protein and histone deacetylation (S4C Fig), suggesting an epigenetic mechanism underlying telomere folding regulation during senescence. The proteins Sir2, Sin3 and Set2 were of particular interest as they antagonize histone acetylation and were reported to be involved in telomere silencing or the regulation of telomeric chromatin (57–64). We tested the identified chromatin modifiers in a Telo-3C assay and found that indeed the deletions of *SIR2, SIN3* and *SET2* were telomere folding defective (Fig 4D). We further confirmed the downregulation of Sir2, Sin3 and Set2 during senescence (*tlc1*) as observed by mass spectrometry (Fig 4A) through western blotting (Fig 4E). In pre-senescent cells and post-senescent survivors, protein levels did not differ from those of telomerase positive wild type cells (Fig 4E). Of note, we observed a modified Sir2 species in senescent cells suggesting a specific regulation of Sir2’s activity, localization or stability during the senescence program (Fig 4E). Telomere length of these mutants is similar to wt length or in the case of *sin3,* only slightly shorter (Fig 4F, S5A Fig). This further supports the notion that telomere length and the state of telomere folding are uncoupled from each other.

**Figure 4:**
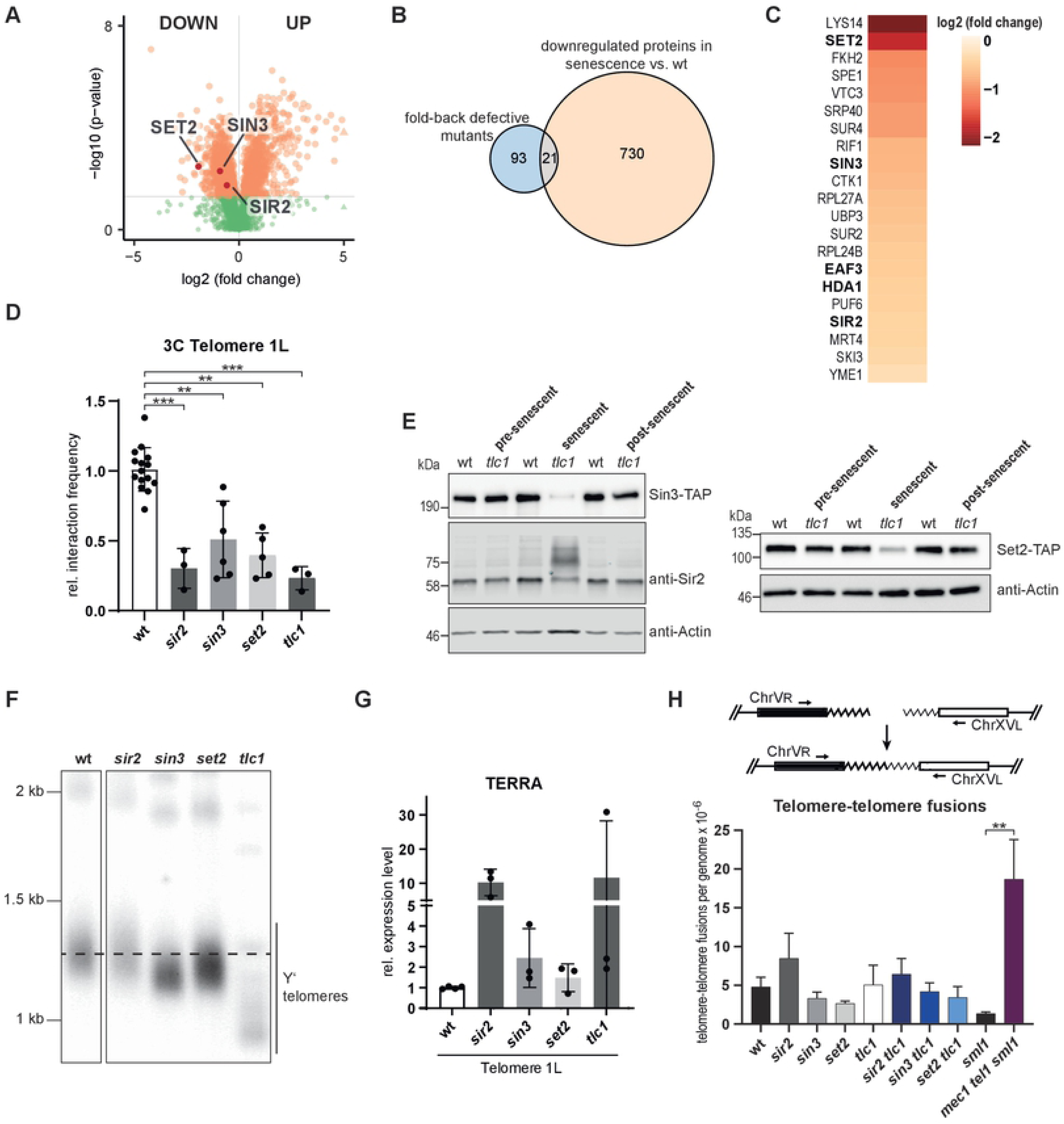
Loss of Sir2, Sin3 and Set2 contribute to telomere opening in senescence. **A.** Volcano plot for up- and downregulated proteins when comparing whole protein extracts from senescent cells (*tlc1*) and extracts from wt cells using label-free quantitative proteomics. Significantly changed proteins are shown in orange (p-value ≤ 0.05). **B.** Venn diagram of genes/proteins overlapping between downregulated proteins in senescent cells (*tlc1*) and mutants defective for telomere folding as described in Poschke et al. 2012. *sir2* was included in the list of fold-back defective mutants. **C.** Heatmap of the 21 overlapping candidates from (B) and the log2 of their fold change during senescence as compared to wt. Histone deacetylases are shown in bold letters. **D.** Telo-3C analysis in *sir2, sin3*, *set2* and *tlc1* (approx. PD40) mutants using primers P5 and P6. The interaction frequency was normalized to an intergenic PCR product. The rel. interaction frequency of wt cells was set to 1. Mean +/- SD of 3-6 independent experiments. Technical replicates were performed. For *sir2* and *tlc1* mutants adjusted p-values were obtained from one-way ANOVA with Dunnetts multiple comparison test. For *sin3* and *set2* mutants adjusted p-values were obtained from unpaired t-tests. (*p ≤ 0.05, **p ≤ 0.01, ***p ≤ 0.001, n.s., not significant) **E.** Western blot for TAP-tagged Sin3 and Set2, as well as Sir2 levels in pre-senescent (approx. PD30), senescent (approx. PD45) and post-senescent cells (approx. PD65). Sir2 and the respective actin loading control were visualized with fluorescent antibodies. All other western blots were developed using HRP-coupled antibodies. **F.** Southern blot for all telomeres using a (TG)n-repeat containing vector fragment labeled with ^32^P. Dashed line indicates the wt telomere length. Superfluous lanes were cropped. The complete blot is shown in S5 Figure. **G.** TERRA levels in in *sir2, sin3*, *set2* and *tlc1* (approx. PD40) mutants. TERRA levels were normalized to Actin RNA levels. TERRA levels of wt cells were set to 1. TERRA levels of the depicted mutants are not significantly different from wt. Mean +/- SD of 3 independent experiments. Technical replicates were performed for the wt. Adjusted p-values were obtained from one-way ANOVA with Dunnetts multiple comparison test. **H.** Scheme of the PCR assay to follow the accumulation of telomere-telomere fusions (T-TFs) (Top). T-TF frequency of the indicated strains as determined by quantitative PCR from liquid cultures, (approx. PD30). Mean +/- SEM of 4-6 independent samples. P-value was obtained from Mann-Whitney test (**p ≤ 0.01).

We estimated the telomere silencing status by measuring levels of the long non-coding RNA (lncRNA) TERRA from telomere 1L. TERRA is a lncRNA transcribed from subtelomeric promoters which are under the influence of local chromatin state (65). As expected, subtelomeres of cells lacking *SIR2* were desilenced and TERRA levels were approx. 10x higher than wt (Fig 4G). As previously reported the increase in TERRA was heterogeneous in *tlc1* cells, due to the stochastic appearance of critically short telomeres (Fig 4G) (66). TERRA levels were also slightly increased in *sin3* and *set2* mutants (Fig 4G). Therefore, the state of telomere folding may have effects on TERRA levels in the cell.

We show here, that Sir2, Sin3 and Set2 are involved in the regulation of telomere folding. The levels of these histone modifiers are downregulated during senescence, which may contribute to the open state during this process. Interestingly, Sir2, Sin3 and Set2 protein levels were not downregulated in a folding-proficient *tel1* mutant with very short telomeres (S5B Fig, Fig 2D, E, F). This observation supports our hypothesis that telomere folding is reduced due to a feature of the senescence program, which may include chromatin alterations through downregulation of chromatin modifiers, but is not caused solely by short telomeres *per se*.

We were wondering what the functional consequences of reduced telomere folding are and if the opening of the telomere structure might lead to a higher incidence of non-homologous end joining (NHEJ) between chromosome ends. We addressed how frequently telomere-telomere fusions (T-TFs) occur in *sir2, sin3* and *set2* mutants by quantitative PCR for T-TFs between chromosomes V and XV (39, 40) (Fig 4H). As expected *mec1 tel1 sml1* cells accumulated T-TFs as previously reported (37–40) (Fig 4H). In contrast, the telomeres of *sir2, sin3* and *set2* mutants as well as those of telomerase negative cells (*tlc1*), did not present an increased incidence of T-TFs. Also the double mutants of *sir2, sin3* and *set2* in combination with loss of *TLC1* showed similar T-TFs frequencies as wt cells (Fig 4H). Therefore, decreased folding does not render telomeres more susceptible to NHEJ.

## Discussion

In this study we adapted the 3C methodology to detect and quantify the interaction of an unmodified yeast telomere with adjacent regions on the juxtaposed subtelomere. Interestingly, interactions between telomere 1L and its subtelomere occurred much more frequently (or stably) than internal interactions (non-telomeric regions) on the same chromosome. We can envisage different explanations for this result. The most straightforward being that DNA ends have a greater mobility as compared to constrained internal loci and hence experience more frequent interactions with nearby loci. Indeed, the interaction frequency of telomere 1L gradually decreases along the chromosome arm towards the centromere, clearly indicating a proximity effect. Alternatively, but not mutually exclusive, the existence of a specific heterochromatin domain at the end of the chromosome may favor the formation of a folded-back structure. This is supported by the genetic dependence of telomere folding on chromatin modifiers such as Sir2, Sin3 and Set2, all of which promote histone deacetylation and hence a more heterochromatic environment. Although Telo-3C can confirm that such telomere-subtelomere interactions are occurring, we cannot infer the stability of the interactions, nor can we gain insights into the structural features of the fold-back. Indeed, the fold-back may actually resemble a lariat or loop with a single touch down point or it may really fold back onto itself and have multiple touching points. We also cannot rule out that the heterochromatic domain at telomeres may exist in the shape of a condensed “knot” rather than the classical fold-back model, whereby interactions within the “knot” occur more frequently and more stably than outside of this structure. It is also tempting to speculate which factors or chromatin landscape might define the borders of this heterochromatin domain. One possibility is that the boarders are defined by a histone mark transition zone, as suggested for an extended silencing domain at subtelomeres (67), mainly concerning H3K79 methylation which is set by the histone methyltransferase Dot1.

Surprisingly, telomere folding is not affected when any of the SMC complexes are depleted. This suggests, that the telomere folding mechanism is independent from the loop extrusion mechanism of cohesin, as well as being distinct from condensin-mediated chromatin condensation. We also observed, that the deletion of the telomeric proteins Rif1 and Rif2, which results in telomere overelongation (4, 5), does not reduce the frequency of telomere folding, as detected by Telo-3C. Moreover, the loss of Ku70, Tel1, Upf1 and Mre11, which leads to extensive telomere shortening, also does not affect the ability of telomeres to fold back. Therefore, telomere length is not a determining factor with respect to telomere structure. The successful telomere folding in *rif1* mutants also suggests that telomere folding is likely different from chromatin architecture forming around a double strand break. It was recently described in human cells that 53BP1 and RIF1 form distinct nanodomains around a double strand break (68). In this case, RIF1 – together with cohesin - was proposed to localize to the boundaries of these domains and stabilize chromatin topology.

Interestingly, we observed decreased telomere folding in cells depleted for the DNA damage checkpoint kinase Rad53. The absence of either one of the upstream kinases, Mec1 (ATR) or Tel1 (ATM), however, did not restrain the proper establishment of telomere folding, suggesting a redundant role for these kinases upstream of Rad53. Alternatively, Rad53 might promote telomere folding in a Mec1 and Tel1 independent manner. That the DNA damage checkpoint factors are important for proper telomere structure is congruent with the finding that in human cells, both ATM and ATR transiently localize to telomeres during the late S and G2 phases of the cell cycle, and are activated there (41, 42). This transient activation of the checkpoint at telomeres might not only be required for completing replication of telomeres but also for processing telomeric overhangs. Additionally, Verdun and Karlseder showed that the checkpoint activation then in turn transiently recruits the homology directed repair machinery, which likely assists in t-loop formation (41). Accordingly, we found that by deleting both of the HDR factors, *RAD51* and *RAD52,* yeast cells were not able to establish a closed state. Taken together, these genetic data strongly suggest that a certain heterochromatic state, that is established by histone deacetylases, together with DDR and HDR factors, promote a fold-back structure (Fig 5A, top) that may be analogous to the “closed state” telomere proposed in human cells. The fact that Rad52 and Rad51 are implemented suggests a strand invasion event into a homologous sequence, which would further the resemblance to mammalian t-loops.

**Figure 5:**
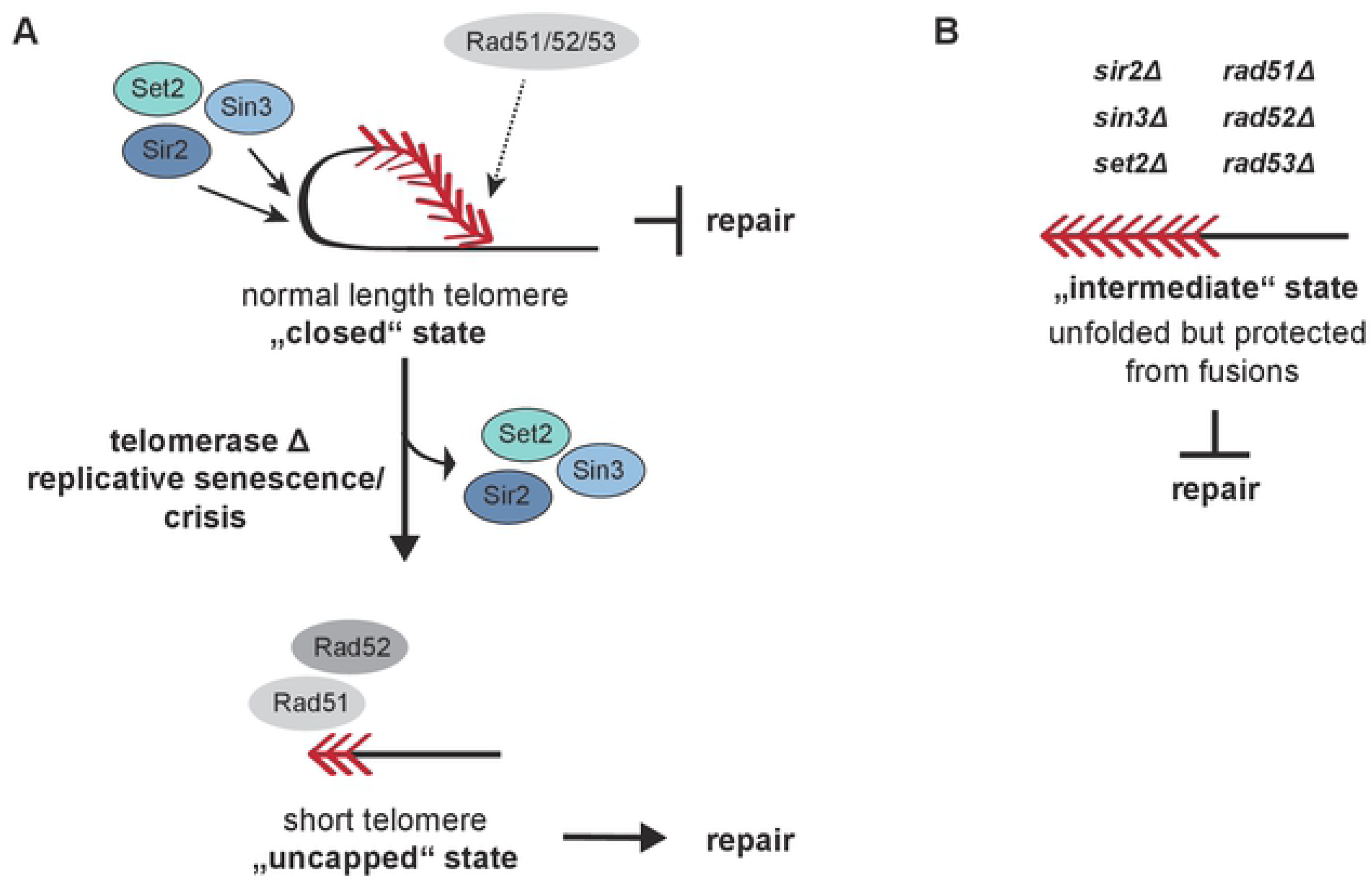
Chromatin modifiers and recombination factors promote a telomere fold-back structure. **A.** Telomere folding is promoted by chromatin modifiers, the checkpoint kinase Rad53 and the recombinases, Rad51 and Rad52. These closed state telomeres are not subject to DNA repair events. When cells enter crisis the loss of chromatin modifiers contributes to the maintenance of an open state. Short and uncapped telomeres in the open state are processed and associate with recombination factors. Due to the absence of Sin3, Set2 and Sir2, a fold-back cannot be established and hence the telomeres are channeled into homology directed repair. **B.** The opening of functional telomeres does not result in telomere repair and telomeres remain capped. This thereby resembles the intermediate state that has previously been described in human cells. (Red arrowheads represent telomeric repeats)

To gain insight into telomere structure during replicative senescence, we used telomerase negative yeast cells (*tlc1*) as a model for this process. During this time course, we found that telomeres transition from a closed, to an open state (unfolded), when cells enter replicative senescence and crisis. Since the opened telomere structure is not due to short telomere length there is likely another feature of the senescence program which is responsible for the opening. This notion is also supported by the observation that telomere folding is decreased after MMS treatment, which was shown to elicit a similar gene expression response as replicative senescence in budding yeast (69). In both cases, cells activate a DNA damage response and a stress response in addition to an altered metabolic program (69). We identified the histone modifiers Sir2, Sin3 and Set2 as potential regulators of telomere folding during senescence. We propose that their reduced protein levels during senescence contribute to ensuring an open state telomere. Sir2, Sin3 and Set2 all antagonize histone acetylation and were reported to be involved in telomere silencing or the regulation of the telomeric chromatin state. The integral role of the SIR complex at telomeres in terms of gene silencing through histone deacetylation, referred to as the telomere position effect (TPE) is well characterized (57,59,70). Sin3 is part of the Rpd3 histone deacetylase complex and is involved in the regulation of silencing on a genome-wide level but also in TPE (60–63). The SET domain containing protein Set2 correlates with histone deacetylation because it methylates histone H3K36 which leads to recruitment of the Rpd3 histone deacetylase complex (Keogh et al. 2005). Sir2 levels did not drastically decrease during replicative senescence, however, we observed an uncharacterized modification of the protein. Sir2 was previously shown to be SUMOylated during mother cell aging in yeast, which leads to its relocalization from telomeres to the nucleolus (71). Therefore, we hypothesize, that during senescence Sir2 may also get redistributed from telomeres to the rDNA locus, contributing to the decreased telomere folding. Accordingly, Sir2 was shown to relocalize from telomeres to other genomic loci under various types of DNA damage (72), which could account for the MMS-induced telomere opening. Interestingly, also Rap1, the direct telomere binding protein, was also shown to be redistributed during senescence (73). This might potentially contribute to decreased SIR binding at telomeres during senescence, since the recruitment of the SIR proteins to telomeres depends on the presence of Rap1. Additionally, it is evident that histone levels themselves drastically decrease during senescence in yeast and human cells (73, 74) and that histone synthesis is sensitive to DNA damage signaling (74). This might also point towards a mechanism by which reduced nucleosome occupancy at subtelomeres contributes to the opening of telomere folding during senescence.

In mutants defective for telomere folding we did not observe an increased incidence of telomere-telomere fusions. Also when telomerase was deleted in addition to the histone modifiers Sir2, Sin3 and Set2, telomere-telomere fusion did not increase. This suggests that the non-homologous end joining repair pathway cannot act on unfolded telomeres and that they remain protected from fusions (Fig 5B). Open, but non-fusogenic telomeres in yeast may be similar to intermediate state telomeres that have been described in human cells as part of the three-state model (75, 76). In this model, telomeres exist in a “closed”, “intermediate” or “uncapped” state. In the fully capped, closed state, the presence of a t-loop was suggested to prevent a DNA damage response that would elicit repair activities at telomeres (as in Fig 5A, top). In the “intermediate” state, telomeres activate a DNA damage response likely due to the loss of the protective loop structure at the chromosome end. This state, however, still retains the capacity to suppress fusions which would correspond to the observed yeast telomeres lacking a folded structure but being resistant to NHEJ (Fig 5B). In the “uncapped” state, telomeres are fully dysfunctional, open, and permissive to repair events (Fig 5A, bottom).

In summary, we propose that the presence of the histone modifiers Sir2, Sin3 and Set2, as well as the DDR and HDR factors, is required for the successful establishment of telomere folding (Fig 5). In wt yeast cells, DNA damage checkpoint factors, such as Rad53, and the HDR machinery (including Rad51 and Rad52) might only transiently associate to telomeres. Here they may assist establishing a telomere fold-back structure, with potential strand invasion of the 3’ overhang. In this scenario, telomere folding might prevent sister chromatid exchange between telomeres and/or limit excessive resection, which corresponds to a “closed” telomere state. However, when either Sir2, Sin3 or Set2 are absent, telomeres do not fold-back efficiently and the telomeric chromatin state is likely altered (Fig 5). We believe, that this situation might represent the “intermediate state” telomeres which lost the folded structure but still suppress telomere fusions.

When telomerase negative cells enter crisis, we observed that levels of Sin3 and Set2 are decreased and that Sir2 becomes modified. We hypothesize that the loss of these histone modifiers contributes to the maintenance of the open state (Fig 5). It is difficult, however, to know if the loss of the histone modifiers is the key event in triggering the opening or if it plays a supportive role in ensuring that the open state is maintained. Nonetheless, the open telomeres in combination with uncapping are subject to processing and associate with Rad52 and Rad51 (32). In this context, the HDR factors are not able to promote telomere folding, due to the absence of the required chromatin modifiers, but rather drive telomere repair through HDR (sister chromatid exchanges and break-induced replication).

Here, we have identified key regulatory factors for establishing telomere folding in budding yeast. Our results point towards the distinct telomeric chromatin environment and the homologous recombination machinery as being major requirements for the folding of yeast telomeres. We provide insight into how the topological reorganization telomeres integrates into the senescence program.

## Material and Methods

### Materials

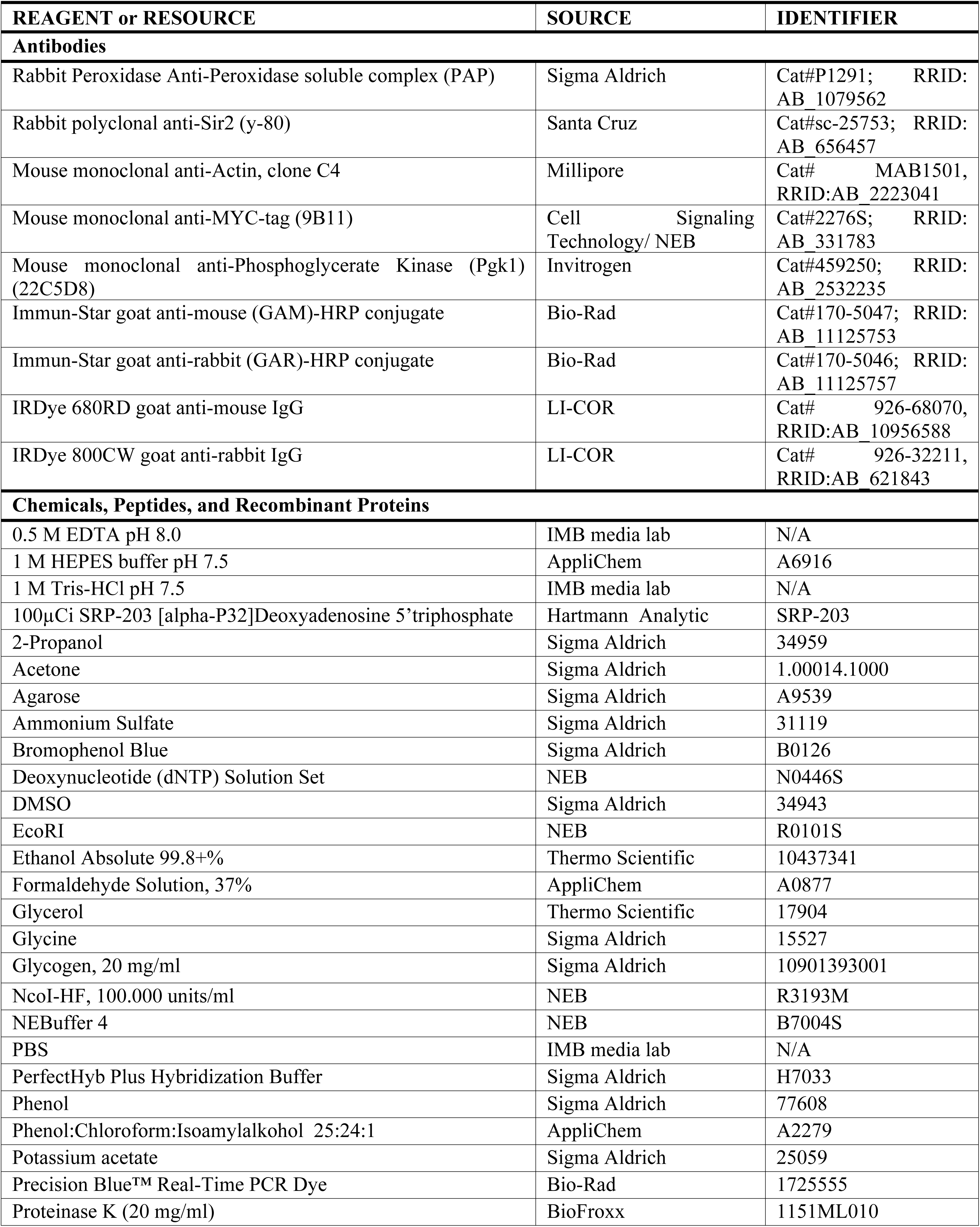
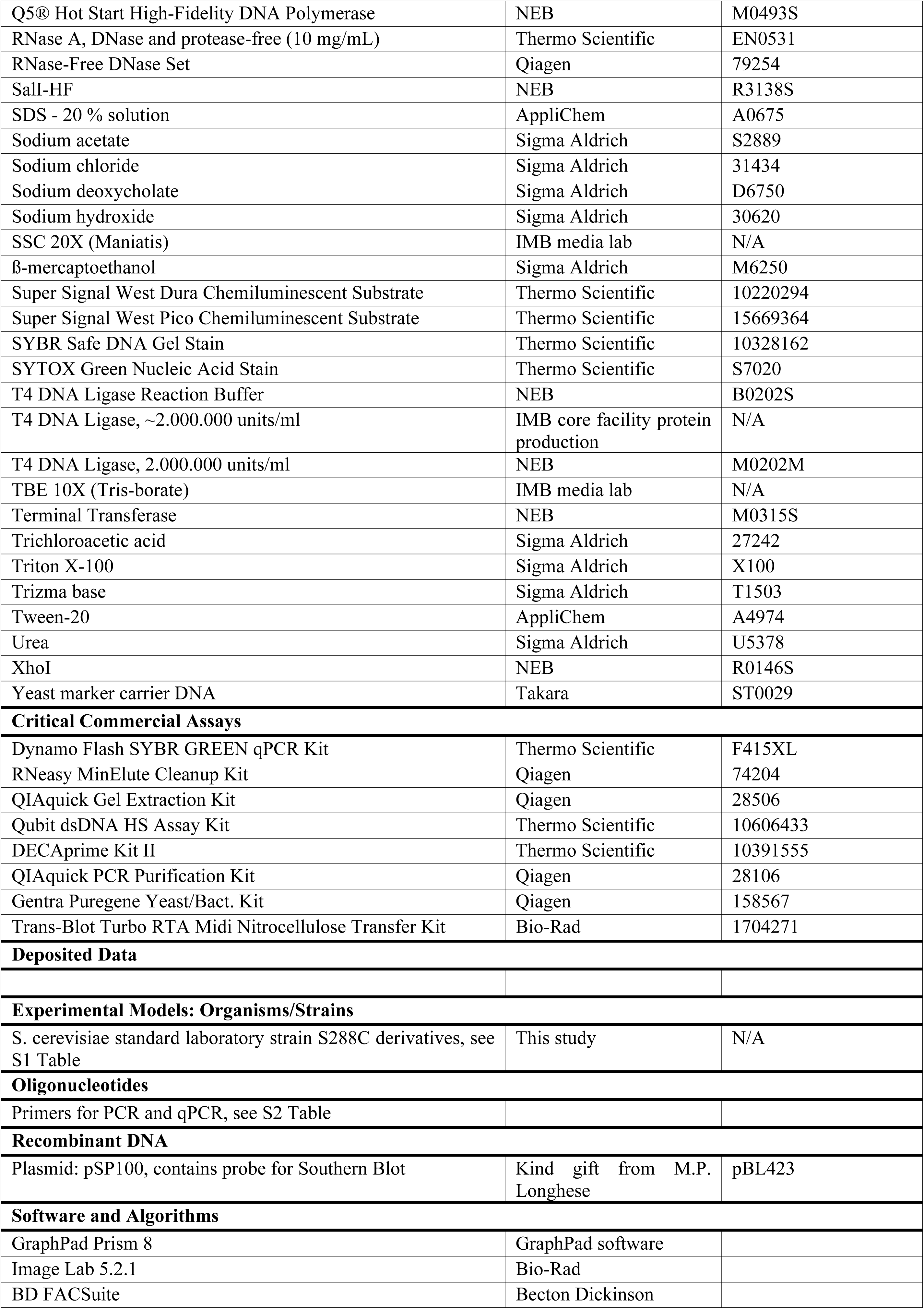
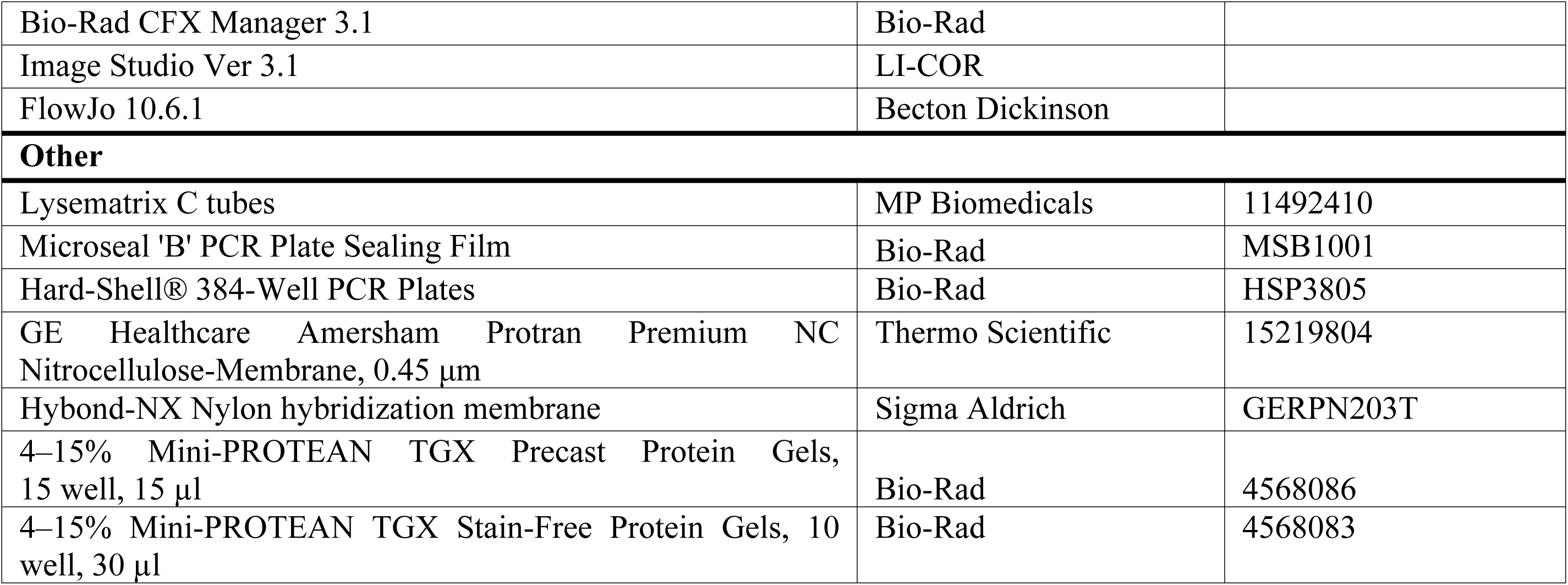

### Methods

#### Yeast strains and plasmids

Strains used in this study are listed in S1 Table, respectively. Strains were grown under standard conditions in YPD (Yeast Peptone Dextrose) or SC without amino acids (Synthetic Complete) at 30 °C if not indicated otherwise.

#### Measurement of telomere length by PCR

Genomic DNA was extracted from 10 ml of exponentially growing cells with the Gentra Puregene Yeast/Bact. Kit (QIAGEN) according to the manufacturer’s protocol. For the C-tailing reaction, 100 ng of genomic DNA were mixed with 0.9 µl of NEB buffer 4 in a final volume of 9 µl. To denature the DNA, the mixture was incubated at 96 °C for 10 min and cooled to 4 °C. 4 Units of terminal transferase (NEB) were added together with dCTP in a final concentration of 0.1 mM. The C-tailing reaction was carried out as follows: 37 °C for 30 min; 65 °C for 10 min; 96 °C for 5 min; hold at 65 °C. 30 µl of pre-heated PCR mix were added to the DNA. The PCR mix contained 4 µl of 2 mM dNTPs, 4 µl of PCR buffer (670 mM Tris-HCl pH 8.8, 160 mM (NH_4_)_2_SO_4_, 70% glycerol, 0.1% Tween-20), 0.4 µl of Q5 hot start polymerase (2 u/µl) and the primers oBL358 and oBL359. The following cycling conditions were used for PCR: 95 °C for 3 min; 45 cycles of 95 °C for 30 s, 63 °C for 15 s and 72 °C for 20 sec; 72 °C for 5 min; hold at 4 °C. PCR products were separated on an 1.8 % agarose gel containing 1X SYBR Safe DNA gel stain and imaged on the Bio-Rad ChemiDoc Touch Imaging System.

#### Flow cytometry analysis for DNA content

0.18 OD_600_ units of cells were collected and the cell pellet was washed with 1 ml of water. Cells were fixed in 70 % ethanol over night at 4°C. After fixation, cells were centrifuged at 13,000 rpm for 5 min and washed in 800 µl 50 mM Tris pH 7.5. RNA was digested in a 50 mM Tris pH 7.5 solution containing 0.25 mg/ml RNaseA for 2 h at 37 °C. 25 µl of 20 mg/ml proteinase K were added and incubated at 50 °C for 2 h. Samples were sonicated for 10 s in a BRANSON sonifier 450 with the following settings: constant mode, 10 sec, power 1. After sonication SYTOX Green was added to a final concentration of 2 µM in 50 mM Tris and analyzed on a BD FACSVerse flow cytometer. 20,000 events were recorded per sample and the data was analyzed with BD FACSuite.

#### Flow cytometry analysis for cell viability

0.68 OD_600_ units of cells were collected and the cell pellet was washed with 1 ml 50 mM Tris pH 7.5. After spinning down the cells for 1 min at 7000 rpm, the pellet was resuspended in 1 ml 50 mM Tris pH 7.5 containing 0.5 μM SYTOX Green. Cells were analyzed on a BD FACSVerse flow cytometer recording 20,000 events. The data was analyzed with FlowJo 10.6.1. As a control sample for dead cells, cells were incubated at 95 °C for 15 min and subjected to the described protocol.

#### RNA extraction and qRT-PCR to measure TERRA levels

RNA extraction, DNase treatment and reverse transcription were performed as described in (66). qPCR analysis was performed by the CFX384 Touch Real-Rime PCR Detection System (Bio-Rad) using DyNAmo Flash SYBR Green (Thermo Scientific).

#### Telo-3C

25 ml of exponentially growing cells were crosslinked with 1.2 % formaldehyde at OD_600_ 0.4 for 20 min at room temperature. The reaction was quenched with 360 mM glycine for 5 min at RT and cells were placed on ice afterwards. The cells were washed twice with cold PBS and the pellet was resuspended in 200 µl FA-lysis buffer (50 mM HEPES-KOH pH 7.5; 140 mM NaCl; 1 mM EDTA pH 8.0; 1 % Triton X-100). Cell lysis was performed in Lysematrix C tubes (MP Biomedicals) using the FastPrep (MP Biomedicals; 2x 30 s with 1 min on ice in between runs; 6.5 M/s). Cell extracts were recovered in FA-lysis buffer containing 0.1 % sodium deoxycholate (SOD) and centrifuged for 15 min at 4°C, 13.000 rpm. The pellet was washed with FA-lysis buffer and resuspended in 200 ml of 20 mM Tris-HCl pH 7.5. 60 µg of this 3C-extract were digested with 100 units NcoI-HF (NEB) over night. The digestion reaction was stopped by incubating the samples at 65 °C for 20 min in the presence of 1 % SDS. The SDS was sequestered with 75 µl of 10 % Triton X-100 and 570 µl of 20 mM Tris-HCl pH 7.5 were added. The chromatin was pelleted for 15 min at 4 °C, 13.000 rpm, and resuspended in 750 µl ligation mixture containing 200 units T4 DNA ligase (NEB/ IMB core facility protein production), 1X T4-Ligation buffer (NEB) and Tris-HCl pH 7.5. After ligation at RT for 5 h, RNA was digested with 20 µg RNase A and crosslinks were reversed with 7.5 µl Proteinase K and 3.5 µl of 20 % SDS at 65 °C over night. To isolate the DNA, the 3C reaction was mixed with 750 µl Phenol:Chloroform:Isoamylalcohol and the DNA was precipitated from the aqueous phase by adding 2 µl glycogen, 75 µl of 3 M KAc and 2 ml of 100 % Ethanol. The DNA pellet was washed with 70 % Ethanol and resuspended in 40 µl ddH_2_O. The DNA concentration was determined with Qubit HS DNA reagent (Thermo Scientific). 1.5 ng of DNA were used per qPCR reaction, performed with CFX384 Touch Real-Rime PCR Detection System (Bio-Rad) and Dynamo Flash SYBR GREEN (Thermo Scientific). The Telo-3C interaction frequency was normalized to an intergenic PCR product (primers oRS39 and oRS40).

#### Protein extraction, SDS-PAGE and western blot

Proteins were extracted from 2 OD_600_ units of yeast cells as described in (66). Protein extracts were loaded on 4 – 15 % precast SDS-PAGE gels (Bio-Rad). Proteins were blotted on a nitrocellulose membrane with the Trans-Blot Turbo Transfer System (Bio-Rad) using the high molecular weight program. The membrane was blocked for 1 h with 5 % skim milk in PBST and primary antibodies were used in the following dilutions in 5 % skim milk in PBST: PAP (Sigma Aldrich) at 1:3000; anti-Sir2 (Santa Cruz) at 1:1000; anti-Actin (Millipore) at 1:2000; anti-MYC-tag (Cell Signaling Technology/NEB) at 1:1000; anti-Pgk1 (Invitrogen) at 1:20,000. The membranes were incubated with the primary antibodies overnight at 4°C. After four washing steps with PBST the respective secondary antibodies were added for 1 h at RT: goat anti-mouse-HRP conjugate (Bio-Rad) at 1:3000; goat anti-rabbit-HRP conjugate (Bio-Rad) at 1:3000; IRDye 680RD goat anti-mouse IgG (LI-COR) at 1:10,000; IRDye 800CW goat anti-rabbit IgG (LI-COR) at 1:10,000. Before detection, membranes were washed three time in PBST and once in PBS.

Membranes with HRP-coupled secondary antibodies were developed using the Super Signal West Pico Chemiluminescent Substrate (Thermo Scientific) or the Super Signal West Dura Extended Duration Substrate (Thermo Scientific) on the Bio-Rad ChemiDoc Touch Imaging System.

Membranes with secondary antibodies coupled to IRDyes were detected on the Odyssey CLx 9140 Imaging System (LI-COR) and processed via the Image studio software (LI-COR).

#### Southern blot

Genomic DNA (5 µg) was digested with the respective restriction enzyme for 5 h at 37 °C (XhoI for all telomeres; SalI for telomere 1L). The digested DNA was separated on a 0.8 % agarose gel overnight at 50 V. DNA in the gel was denatured for 1 h (0.4 M NaOH, 0.6 M NaCl) and neutralized for 1 h (1 M Trizma Base, 1.5 M NaCl, pH 7.4). DNA was transferred to a nylon membrane (Hybond NX, GE Healthcare) via capillary transfer in 10X SSC overnight and crosslinked to the membrane with UV light (auto X-link, UV Stratalinker 2400, Stratagene). The membrane was pre hybridized for 5 h at 55 °C in hybridization solution (PerfectHyb™ Plus Hybridization Buffer, Sigma Aldrich). Hybridization of the respective radio-labeled probe was carried out overnight at 55 °C. The membrane was washed twice in 2X SSC with 0.1 % SDS for 5 min and twice in 0.5X SSC with 0.1 % SDS for 20 min. All washing steps were performed at 55 °C. The membrane was exposed to a phosphoimager screen and the signal was detected via Typhoon FLA 9500 (GE Healthcare).

#### Probes for Southern blot

The telomere-specific probe was obtained by digestion of pBL423 with EcoRI followed by gel extraction using the QIAquick Gel Extraction Kit (Qiagen). Plasmid pBL423 was a kind gift from M.P. Longhese (pSP100). The probe for subtelomere 1L was generated by PCR with oligos oSM1 and oSM2 on genomic DNA of wt *S. cerevisiae* cells. The PCR product was separated on a 1 % agarose gel and purified using the QIAquick Gel Extraction Kit (Qiagen). 60 ng of each probe were labeled with dATP [α-32P] (DECAprime kit II; Thermo Scientific) by random priming according to the manufacturer’s instructions.

#### Senescence curve

Liquid senescence assays were performed as described in Balk et al. 2013. Briefly, the OD_600_ of every culture was measured every 24 h and cultures were diluted back to an OD_600_ 0.01 subsequently. All analysis during the senescence curve were performed on exponentially growing cultures, starting from the saturated culture after 24 h.

#### MS/MS

1 OD_600_ unit of exponentially growing cells was resuspended in 1X LDS buffer supplemented with 100 mM DTT. Cells were heated for 10 min at 95 °C and separated on a 4-12 % NuPAGE Bis –Tris precasted PAGE gel (Thermo Scientific). Samples were run at 180 V for 10 min and processed by in-gel digestion (78). Briefly, samples were reduced in reduction buffer (10 mM DTT in 50 mM ammonium bicarbonate (ABC) buffer) for 1 h at 56 °C and alkylated in alkylation buffer (50 mM iodoacetamide (IAA) in 50 mM ABC buffer) for 45 min in the dark. Proteins were digested with 2 µg Protease LysC (Wako Chemicals) overnight at 37 °C in 50 mM ABC buffer. Digested peptides were desalted on a C18 StageTip as described (79) and analyzed by nanoflow liquid chromatography on an EASY-nLC 1000 system (Thermo Scientific) coupled to a Q Exactive Plus mass spectrometer (Thermo Scientific). The peptides were separated on a self-packed reverse phase capillary (75 µm diameter, 25 cm length packed with C18 beads of 1.9 µm (Dr Maisch GmbH). The capillary was clamped on an electrospray ion source (Nanospray Flex™, Thermo Scientific). A 90 min gradient starting from 2 % - 60 % gradient acetonitrile in 0.1 % formic acid was used at a flow of 225 nl/min. Data was collected in data-dependent acquisition mode with one MS full scan followed by up to 10 MS/MS scan with HCD fragmentation.

#### MS data processing and bioinformatics analysis

MS raw files were processed using the MaxQuant software (version 1.5.2.8) and the ENSEMBL *S. cerevisiae* protein database (Saccharomyces_cerevisiae.R64-1-1.24). LFQ quantitation and match between run options were activated. MaxQuant output files were analyzed using an in-house R script. Known contaminants, reverse hits and protein groups only identified by site modification were excluded. Identified protein groups (minimum 2 peptides, 1 of them unique) were further filtered to a minimum of 2 quantification events per experiment. Missing values were imputed using a downshifted and compressed beta distribution within the 0.001 and 0.015 percentile of the measured values for each replicate individually. The LFQ intensities were log_2_ transformed and a two sample Welch t-test was performed. Volcano plots were generated by plotting -log_10_(p-values) and fold changes. The threshold line for enriched proteins is defined with p-value = 0.05. Gene ontology analysis were performed with the PantherDB.org (80) overrepresentation Test (Release 20190711) with the annotation database released on 20190703. Fisher’s exact test followed by FDR correction was applied to calculate the p-values. Heatmap for enriched proteins were generated using the “pheatmap” (version 1.0.12) or “ggplot2” (version 3.2.1) package in R. Additionally the Venn diagram was generated using the “eulerr” (version 5.1.0) package in R (version 3.5.1).

#### Telomere-telomere fusions analysis

Telomere-telomere fusions (T-TFs) were analyzed by semiquantitative and quantitative PCR analyses as reported (39), with the modifications described in (40). Briefly, ∼100 ng of *Sau3*A-treated genomic DNA extracted by standard protocols from asynchronous cultures was PCR-amplified for semiquantitative analyses using a primer from the X element of chromosome XV-L and a primer from the Y’ element of chromosome V-R. A DNA fragment from HIS4 was PCR-amplified as input control. T-TFs and HIS4 were PCR-amplified using 35-40 or 20 cycles, respectively, under the conditions previously reported, except for the annealing temperature (60 °C) and the extension time (30 s). Quantitative analyses were performed by real-time PCR by using the same amount of *Sau3*A-treated genomic DNA, the oligonucleotides used for semiquantitative analyses and the PCR conditions described previously (Mieczkowski et al, 2003). The frequency of T-TFs per genome was calculated with the formula: T-TFs/genome = 2^−N^ / N = Ct (T-TFs) – Ct (*HIS4*). Prior to applying this formula, the curves representing the increasing amounts of DNA for the two products as a function of the number of PCR cycles were confirmed to be parallel (i.e., the slope of the curve representing the log of the input amount versus ΔCt was < 0.1).

#### Construction of strains with auxin inducible degrons

Strains carrying auxin-inducible degrons (AID*) for SMC complexes were created as described before (81). Primers for tagging are listed in S2 Table.

The *RAD53-AID*-9MYC* construct was transferred from the strain published by Morawska and Ulrich 2013 into the S288C background. For this purpose, the c-terminal sequence of *RAD53* including the tag was amplified by PCR and integrated into a strain carrying *leu2::AFB2::LEU2* (yKB244).

## Acknowledgements

We thank the Luke lab members for support and discussions, the Media lab, Flow cytometry and Protein production core facilities of the IMB. We thank Alex Orioli, Annika Müller, and Diego Bonetti in the assistance with strain construction. BL’s lab was supported by the Heisenberg Program of the DFG - LU 1709/2-1. Research in FP’s lab was funded by grants from the Spanish government (BFU2015-63689-P).

## Supporting informatio

### Supporting Figure Legends

**S1 Figure:**

**A.** Southern blot for telomere 1L on genomic DNA digested with SalI using a specific PCR product labeled with ^32^P. DNA was stained with Red Safe as a loading control. One representative sample per genotype is shown.

**B.** Flow cytometry histograms for DNA content of wt and *tlc1* cells during the time course shown in (Fig 2A). One representative sample per genotype is shown.

**S2 Figure:**

**A.** Flow cytometry histograms for viability using SYTOX green stain after the depletion of the SMC complex members for 1 h (treatment with 1 mM IAA). As a control for dead cells, wt cells were incubated for 15 min at 95 °C.

**B.** Flow cytometry histograms for viability using SYTOX green stain after the depletion of Rad53 for 2 h (treatment with 1 mM IAA). As a control for dead cells, wt cells were incubated for 15 min at 95 °C.

**S3 Figure:**

List of all significantly up- or downregulated proteins in *tlc1* versus wt cells (p ≤ 0.05) with log2 fold change (blue: upregulated; red: downregulated).

**S4 Figure:**

**A.** Top 20 gene ontology (GO) terms for biological processes of 751 proteins downregulated in senescence compared to wt.

**B.** Top 20 GO terms for biological processes of 678 proteins upregulated in senescence compared to wt.

**C.** Results from gene ontology analysis for molecular function of the 21 overlapping candidate genes from (Fig 4B). Shown are GO terms with p-values below 0.05. *sir2* was included in the list of fold-back defective mutants.

**S5 Figure:**

**A.** Uncropped Southern blot corresponding to Fig 4F.

**B.** Western blot of TAP-tagged Sin3 and Set2, as well as Sir2 levels in *tel1* mutants.

### Supporting Table Legends

**S1 Table:**

Yeast strains used in this study

**S2 Table:**

Primers used in this study

